# Multi-Modality Deep Infarct: Non-invasive identification of infarcted myocardium using composite in-silico-human data learning

**DOI:** 10.1101/2024.05.31.596513

**Authors:** Rana Raza Mehdi, Nikhil Kadivar, Tanmay Mukherjee, Emilio A. Mendiola, Dipan J. Shah, George Karniadakis, Reza Avazmohammadi

## Abstract

Myocardial infarction (MI) continues to be a leading cause of death worldwide. The precise quantification of infarcted tissue is crucial to diagnosis, therapeutic management, and post-MI care. Late gadolinium enhancement-cardiac magnetic resonance (LGE-CMR) is regarded as the gold standard for precise infarct tissue localization in MI patients. A fundamental limitation of LGE-CMR is the invasive intravenous introduction of gadolinium-based contrast agents that present potential high-risk toxicity, particularly for individuals with underlying chronic kidney diseases. Herein, we develop a completely non-invasive methodology that identifies the location and extent of an infarct region in the left ventricle via a machine learning (ML) model using only cardiac strains as inputs. In this transformative approach, we demonstrate the remarkable performance of a multi-fidelity ML model that combines rodent-based in-silico-generated training data (low-fidelity) with very limited patient-specific human data (high-fidelity) in predicting LGE ground truth. Our results offer a new paradigm for developing feasible prognostic tools by augmenting synthetic simulation-based data with very small amounts of in-vivo human data. More broadly, the proposed approach can significantly assist with addressing biomedical challenges in healthcare where human data are limited.

## 1 Introduction

Myocardial infarction (MI), commonly known as a heart attack, is a critical and life-threatening cardiovascular condition that claimed the lives of one million people in 2019 and continues to be a leading cause of death in the United States^1,2^. This acute condition occurs when the blood supply to the cardiac muscle is blocked, leading to the death of the myocardium^3,4^. The long-term consequences of MI can be profound, often leading to chronic cardiac dysfunction and an increased risk of additional subsequent heart attacks^5,6^. Despite improvements in post-infarction survival rates following technological advances^7^, MI often leads to chronic heart failure (HF), which is the leading cause of mortality associated with MI^8,9^. Scar formation in the myocardium following MI significantly impacts cardiac function^10^, with the size and extent of the scar (infarct region) evidently playing an important role in the long-term outcome of MI^5^. Indeed, the precise quantification of infarcted tissue within the whole myocardium has emerged as a crucial part of diagnosis, therapeutic management, and improved post-MI care^11^. Additionally, assessing the extent and location of myocardial infarct remains essential for developing personalized risk stratification and treatment protocols^12^.

Electrocardiogram (ECG) has served as a valuable tool in the detection and prognostic assessment of MI. Various studies have utilized ECG data, leveraging machine learning (ML) algorithms and signal processing techniques, to achieve high detection accuracy. In particular, Dohare et al.^13^ employed support vector machines (SVM) on the Physikalisch-Technische Bundesanstalt (PTB) dataset^14^ and achieved excellent accuracy by expediently selecting clinical features of 12-lead ECG, including P duration, QRS duration, ST-T complex interval, and QT interval. Similarly, as a prominent R wave can indicate MI, Wang et al.^15^ utilized the Pan–Tompkins algorithm^16^ for R-wave peak detection from ECG signals using the same PTB dataset, and subsequently applied ML algorithms such as k-nearest neighbor (KNN), backpropagation neural networks (BPNN), SVM, and random forests (RF). Their findings highlighted the superiority of the RF algorithm in terms of detection accuracy.

Moreover, Baloglu et al.^17^ introduced a deep convolutional neural network (CNN) model to detect MI using ECG data from the PTB dataset and achieved a remarkable accuracy of 99%. Along these lines, several other studies employed ECG signals for MI detection and surpassed human capabilities in terms of accuracy^18–20^. While these advancements showcase the potential of ECG in MI detection, it is crucial to note that ECG primarily identifies the presence or absence of MI but falls short in pinpointing the precise location and extent of myocardial damage. Pinpointing injured myocardium is key in MI prognosis, assisting in the determination of the most effective revascularization approach, assessment of the extent of cardiac damage, and facilitation of accurate evaluation of regional tissue remodeling^21,22^. Among other therapies, precise MI identification is essential for developing potential localized tissue reinforcement therapies, such as hydrogel injection and cardiac patches^23,24^. The utilization of cardiac magnetic resonance (CMR) with late gadolinium enhancement (LGE) is regarded as the gold standard for precise infarct tissue localization in MI patients. While this imaging method provides the most precise delineation of MI regions, it is not without limitations and presents several inherent drawbacks within the clinical setting^25–27^. One notable limitation lies in the risk it imparts upon patients, primarily due to the invasive intravenous introduction of gadolinium-based contrast agents (GCA) throughout the imaging process^25^. This presents a potential high-risk scenario, particularly for individuals with underlying chronic kidney diseases, as the administration of gadolinium-based CA can have fatal consequences in such patients^25^. Moreover, the reliance on CMR imaging for infarct localization undermines the affordability and ubiquity of this procedure, with echocardiography being the safest, most cost-effective, and ubiquitous cardiac imaging modality. Additionally, LGE-CMR introduces non-repeatability and time-consuming factors into the clinical workflow. The process relies on experienced clinicians to detect and manually segment “bright” areas from the LGE-CMR images, a step that is susceptible to considerable variation in interpretation among different observers^27^. This multi-step process, which includes LGE in CMR imaging, manual segmentation, and quantification of segmented areas, not only extends the time needed for diagnosis but may also result in cumulative errors^26^. In light of these limitations, there is a pressing need to develop alternative, less invasive, and more efficient approaches to infarct localization in MI patients, which can address the shortcomings associated with LGE approach and its reliance on CMR, GCA, and manual segmentation. To overcome the limitations associated with LGE-CMR imaging, we propose the use of cardiac strains derived from myocardial deformation analysis as a promising alternative to detect and characterize infarct regions without the use of LGE in imaging. Several studies have investigated the ability of cardiac strains to identify MI. For example, Sengupta et al.^28^ employed speckle-tracking echocardiography to assess circumferential and longitudinal strains, demonstrating their ability to identify MI-induced myocardial dysfunction. Similarly, Wu et al.^29^ utilized CMR tagging to quantify circumferential and radial strains, providing valuable insights in to regional myocardial mechanics in MI patients. Moreover, techniques such as feature tracking, strain-encoded CMR, and displacement encoding with stimulated echoes (DENSE) have been employed to measure various components of cardiac strains, further enhancing the precision of MI characterization^30–32^. While these studies have provided significant insights in to the capacity of cardiac strains as markers to detect infarct myocardium, the need to develop a standardized tool to locate infarct and subcontractile regions in the heart that can be seamlessly integrated into routine clinical use persists.

In this work, we introduce a ML approach to assess the location and extent of MI regions in the left ventricle (LV). The ML model predicts the infarct size and location utilizing circumferential, radial, and longitudinal (CRL) strains, which can be readily extracted from routine CMR images and potentially other cardiac imaging modalities, including echocardiography. We proposed a single-fidelity ML model based on the UNet architecture^33^trained on data from low-fidelity rodent computational cardiac models (RCCMs). This model was trained using CRL strains at end-systole (ES) obtained from RCCM simluations, enabling it to predict the location and extent of MI regions. We then extended the single-fidelity ML model into a composite neural network architecture^34^ that incorporated multi-fidelity data (RCCM + LGE-CMR human data). This multi-fidelity modeling approach was used to enhance the predictive accuracy of the ML model by augmenting it with human LGE-CMR data. Multi-fidelity methods have gained prominence in recent years for their ability to leverage heterogeneous data sources with different fidelity to improve the overall accuracy and efficiency^35–37^. These methods are particularly effective in scenarios where high-fidelity data are very limited and expensive to obtain, while low-fidelity data are abundant and easily affordable. Often, low-fidelity data can provide valuable insights into the trends of high-fidelity data, thus enhancing the predictive accuracy of the multi-fidelity modeling approach when compared to single-fidelity modeling, even with a limited set of high-fidelity data^38–40^. The multi-fidelity framework offers significant advantages in terms of efficiency and effectiveness, as it can model complex correlations between data of varying fidelity^41^. The trained ML model can seamlessly integrate with the CMR image processing pipeline, extracting strains from raw CMR images, which then serve as inputs to the ML model for infarct region prediction. Our findings indicate that the proposed ML model is proficient at learning the complex relationship between cardiac strains and infarct labels, making it a promising tool for infarct region estimation.

## 2 Materials and methods

### Overview

We developed subject-specific finite-element (FE) RCCMs constructed by integrating in-vivo hemodynamic data with ex-vivo mechanical and imaging data obtained from infarcted hearts. The development of the RCCMs is briefly described in the following subsection, and a comprehensive, detailed description of the RCCM development is reported in Mendiola et al.^42^. Moreover, we developed a library comprising 592 hearts characterized by varying infarct sizes and stiffness levels. Extensive simulations were conducted to generate a large diverse dataset used to train and test the proposed ML models. Finally, the trained models were evaluated against human LGE-CMR data to further assess the efficacy and accuracy of our single-fidelity and multi-fidelity approaches.

### 2.1 Rodent cardiovascular computational model simulations

#### 2.1.1 Animal model of myocardial infarction

The procedures performed on the animals in this study followed protocols that adhered to the guidelines approved by the ethical treatment of Institutional Animal Care and Use Committee (IACUC) at the Texas Heart Institute (THI). Four male Wistar-Kyoto (WKY) rats, aged 8 weeks at the beginning of the experiment, were utilized. Anteriobasal infarct was induced through ligation of the left anterior descending artery near the base of the heart. The infarcted rats were sacrificed at four timepoints post-MI (1-, 2-, 3-, and 4 weeks, n=1 at each timepoint). Immediately prior to sacrifice, pressure-volume (P-V) measurements of the LV were collected; subjects in the infarct group were then given GCA in the tail vein. The heart was excised and flushed with phosphate-buffered saline. The ventricles were then filled with an octreotide solution to approximately ED pressure. Heart tissues were then fixed in a 10% formalin solution. High-resolution standard CMR was conducted, in addition to T1-weighted and diffusion tensor imaging (DTI), and employed to develop FE RCCMs, starting with reconstructing the three-dimensional (3-D) biventricular model as described below. In a parallel animal study^42,43^, 24 similar WKY rats (n=6 at each timepoint post-MI) were used for biaxial mechanical testing. These rats provided passive stiffness values used in the <u>“RCCM development”</u> section. The statistical spreads in stiffness values of the healthy and infarct regions for each timepoint were used to provide variations of stiffness (described in <u>“Generation of RCCM library”</u> section) across the four geometries originating from the initial four hearts examined in this study.

#### 2.1.2 Infarct identification and fiber architecture mapping

The 3-D biventricular heart geometries were reconstructed from CMR scans truncated at the valve plane, and each geometry was discretized using tetrahedral elements (Fig.1). The remote and infarct regions of the post-MI heart models were determined by identifying areas of the LGE CMR that showed increased contrast relative to remote myocardium. The infarct region was also evident in DTI scans as a region with no signal or disturbed fibers. The myofiber orientation distribution in the remote region, reconstructed from DTI scans and weighted by principal directions at each voxel, was registered to the corresponding ventricular mesh from the same rodent heart.

**Fig. 1.**
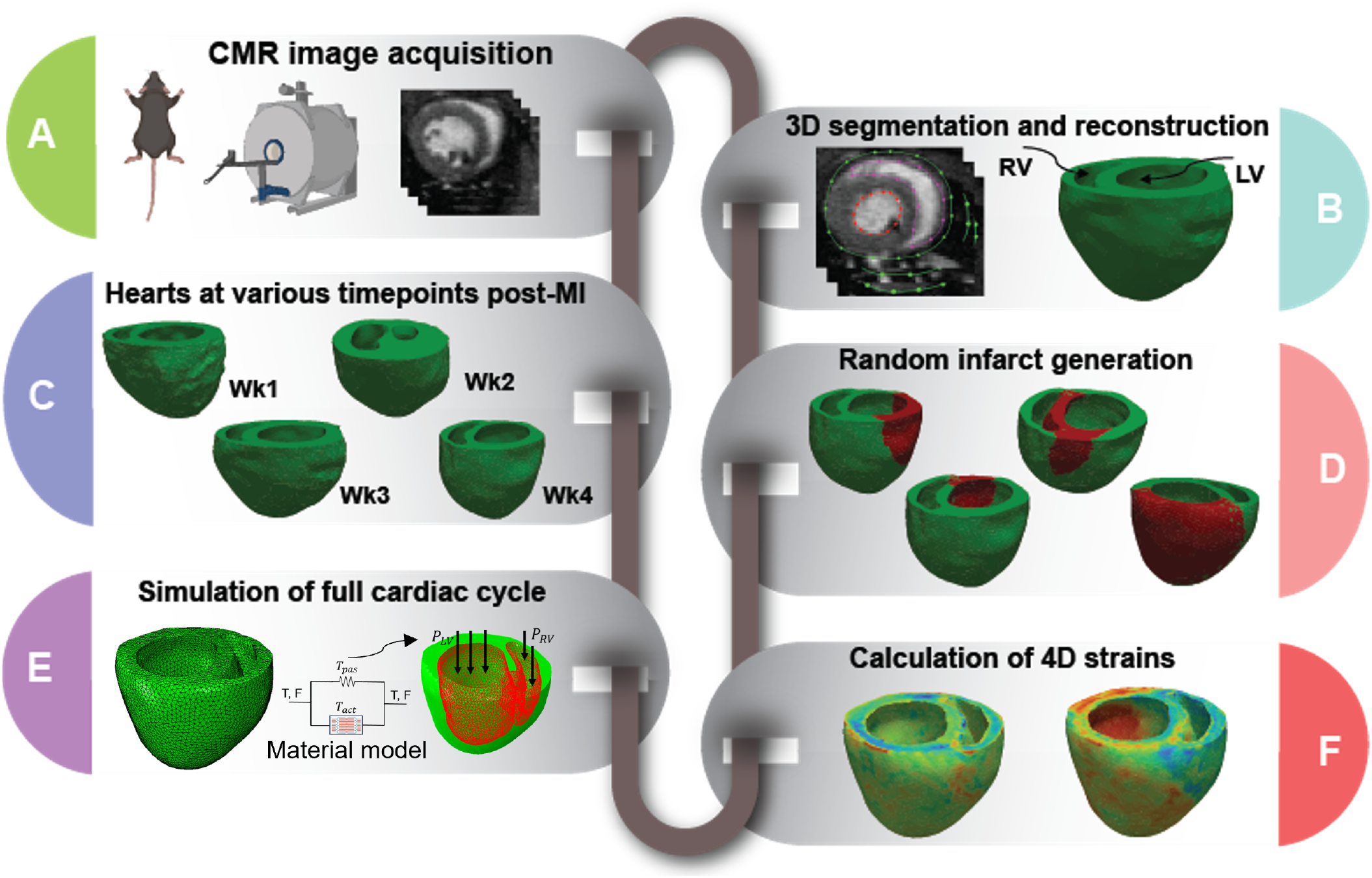
Development of simulated cardiac strains by the use of computational rodent heart models involved reconstruction of 3-D cardiac geometry from cardiac magnetic resonance (CMR) images, creation of a library of finite-element models with random infarct size and location from cardiac geometries at different time points of post-myocardial infarction, and estimation of cardiac strains via finite-element simulation using experimentally derived passive and active properties.

#### 2.1.3 Constitutive model for the myocardium

A transversely isotropic hyperelastic material model (characterized by the myofiber direction) was used to capture both passive and active behaviors of the healthy myocardium through an additive stress decomposition^44^, given by:

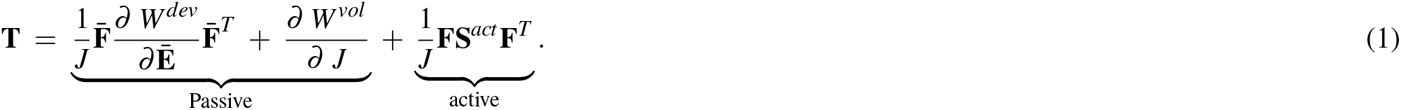

where **T** represents the total Cauchy stress, **F** is the deformation gradient, *J* denotes volumetric changes in deformation, and 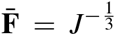 **F** represents the deviatoric part of **F**. The passive stress follows a Fung-type strain energy function *W* (**E**, *J*) given by^45^:

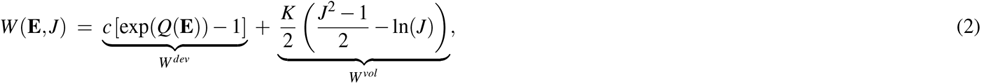

where *W*^*dev*^ and *W*^*vol*^ denote the deviatoric and volumetric components of *W*, and 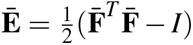 is the Green-Lagrange strain tensor. The quadratic form *Q* is expressed as: 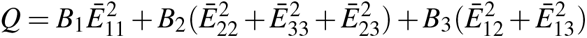 relative to the Cartesian coordinate system *{***e**_1_, **e**_2_, **e**_3_*}*, with **e**_1_ indicating the local fiber direction. Constants *B*_1_, *B*_2_, and *B*_3_ are dimensionless constants representing local anisotropy in the myocardium, *c* is a positive stress-like constant, and *K* is the bulk modulus.

A constitutive equation for the active stress can be written in terms of the second Piola–Kirchhoff active stress tensor as

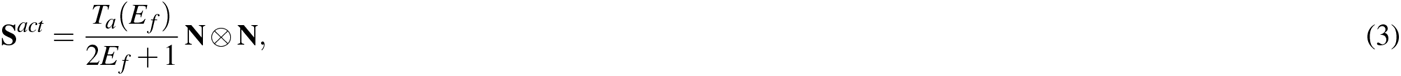

where *T*_*a*_(*E*_*f*_) is a stress-like positive function of the strain in the fiber direction **N** given by *E*_*f*_ = **N** *·* **EN**. We chose the following form for *T*_*a*_(*E*_*f*_)

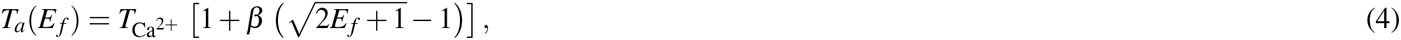

following the Hunter-McCulloch-TerKeurs model^**?**^ for the mechanical behavior of a contractile myocyte.

The passive behavior of the myocardium was characterized by four material parameters. Specifically, distinct material parameters were assigned to healthy and infarcted regions. For the healthy remote myocardium, the parameters *c*_*r*_, 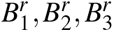 were utilized, while the infarct region employed parameters *c*_*i*_, 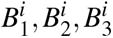. These parameter values were determined through an analytical fitting process of the constitutive model (Eq. 1) to biaxial experimental data^43^. The active force parameter 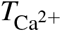 was estimated by reproducing the experimental P-V loop of and minimizing any differences between the simulated and experimental results. The infarct region had no contractile behavior. To conduct forward FE simulations, both passive and active properties were assigned to these geometries. Detailed information regarding the experimental measurements and the corresponding properties obtained from those measurements is described by Mendiola et al.^42^.

#### 2.1.4 Generation of RCCM library

A comprehensive library comprising 592 diverse heart models was developed, building upon the initial 4 models. Random infarct regions were created in the LV in all the reconstructed hearts as described below (Fig.1D). Statistical analysis was conducted to elucidate the correlation between the average strains of infarcts and both the percentage of infarct size in the left ventricle, as well as the stiffness of the left ventricle (Supplementary Fig. 1). A random point in LV was selected, serving as the starting location for each infarct. Next, a random percentage of the total elements, ranging from 5% to 60%, was chosen to determine the infarct size at that particular location. This process was repeated for all the reconstructed hearts, totaling 592 examples, generating a diverse set of heart models with infarcts of varying sizes and locations across the LV. To ensure a non-repetitive and evenly distributed selection of infarct locations and sizes, Latin hypercube sampling (LHS) methodology was employed, which enabled systematic and uniform coverage of the parameter space, thus mitigating any potential bias in the infarct generation process. Inverse optimization was used to determine stiffness values in the infarct regions using biaxial stress-strain data for all post-MI timepoints (n=6 per timepoint)^42,43^. The standard deviations of the stiffness values did not exceed 30% (relative to the mean values) for any post-MI timepoint. Therefore, we applied a Latin hypercube sampling approach with 30% upper and lower bounds (relative to the mean value) to generate the stiffness values. This way, we generated distinct stiffness values for each example. A forward simulation was performed with each unique cardiac model in the library, simulating a full cardiac cycle. Following the forward simulation, cardiac CRL strains were obtained for the entire heart at end-systole (ES) (Fig.1).

### 2.2 Preprocessing of ML inputs

#### 2.2.1 American Heart Association-based representation of strains

The focus of this study was to locate infarcts within the LV. For this purpose, the cardiac CRL strains of LV were obtained at the base, mid, apical, and apex short-axes, representing four common short-axis sections in routine CMR scans. Next, the strain data was translated into standard American Heart Association (AHA) bullseye maps (Fig. 2) showcasing the spatial distribution of the strains across the LV. This conversion procedure was executed for each CRL strain, thereby generating three distinct AHA-based bullseye maps per simulation. The strain maps were stored in images to facilitate further analysis and streamline the input features for ML algorithms. This image-based representation ensured consistency in the input features and allowed for easy preprocessing prior to feeding images into the ML models. By adopting this approach, the strains were effectively visualized in terms of pixels, enabling effective data preprocessing techniques and smooth and cohesive integration with ML models.

**Fig. 2.**
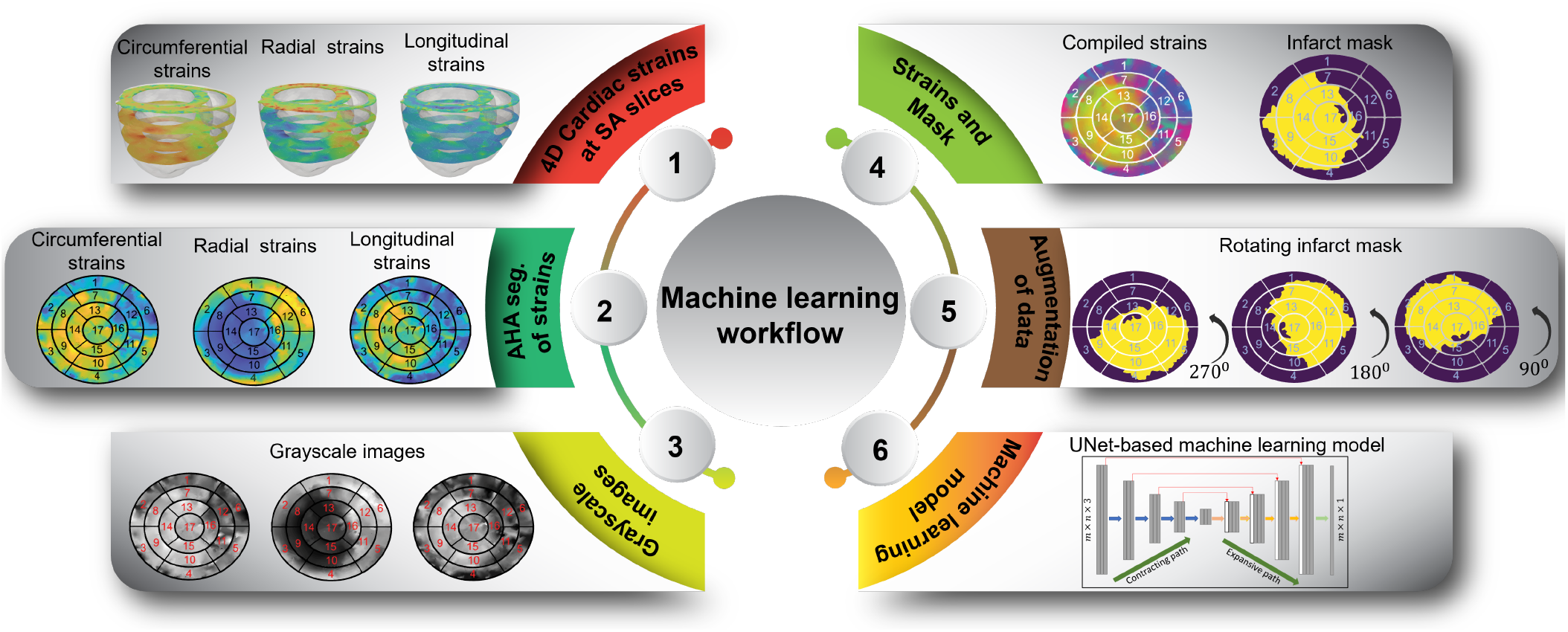
Preprocessing steps: The cardiac circumferential, radial, and longitudinal (CRL) strains of the left ventricle (LV) were acquired at the base, mid, apical, and apex levels. These strain values were then mapped onto standard American Heart Association (AHA) bullseye maps and subsequently transformed into greyscale images. The integration of these three sets of images, together with their corresponding infarct masks, constituted the dataset employed for machine learning (ML) training after the application of data augmentation techniques.

#### 2.2.2 Conversion to greyscale images and normalization

The bullseye maps representing CRL strains served as preliminary inputs to our ML models (Fig. 2). Appropriate preprocessing steps were deemed necessary to optimize the efficiency of using three separate bullseye maps as inputs. In particular, recognizing that the bullseye maps could be generated using any color scheme while maintaining consistent color intensity throughout, we performed a transformation to convert the RGB images representing each type of CRL strain into greyscale images, effectively transitioning from three-channel images to one-channel images. Subsequently, we combined these greyscale CRL images into a single RGB image, wherein the red (R), green (G), and blue (B) channels represented the greyscale image of circumferential, radial, and longitudinal strains, respectively. Having unified the greyscale CRL strain images into a multi-channel representation, we proceeded with an essential normalization step. This normalization process standardized the pixel values across the entire image, thereby promoting smoother convergence during the training of our ML models. By applying this series of preprocessing steps, we aimed to enhance the efficacy of our ML algorithms. The transition to greyscale images and pixel normalization not only accelerated training convergence but also facilitated more meaningful and efficient utilization of the bullseye maps as input features for our ML models. Ultimately, these preprocessing measures played a pivotal role in enhancing the overall performance and accuracy of our ML-driven CRL strain prediction framework.

#### 2.2.3 Data augmentation

After the necessary preprocessing steps, our methodology involves data augmentation to enhance the diversity and robustness of the training process. To this end, we rotated each channel of the image by 90, 180, and 270 degrees as part of a rotation augmentation procedure. This process resulted in a fourfold increase in the number of images within the training dataset. Next, the augmented dataset was partitioned into distinct sets for training and validation purposes, facilitating the training of the ML model. A detailed breakdown of the exact number of images at each stage of the data splitting and augmentation process is presented in Fig. 5. Using the fixed rotation strategy instead of random rotation, we avoided the misalignment of the images and ensured consistency across the dataset. In addition, we applied the rotation procedure after resizing the images into equal width and height for further consistency. This procedure is expected to increase the adaptability of the model by enhancing its capacity for generalization.

### 2.3 Machine learning models

Our aim was to develop and apply ML models to identify infarct regions from LV strain data represented as three-channel images, where each channel corresponds to a distinct directional strain. We began with four encoder-decoder-based designs, namely, UNet, attention UNet, dense UNet, and residual attention UNet, which are known for their precision in segmenting based on spatial variations between pixels in the image^46^. These architectures use the encoder-decoder structures to extract significant characteristics from the images, ensuring that they do not lose any important spatial information. Given this ability, applying these ML models to our strain data, they are expected to identify subtle changes in strain patterns that may indicate the presence of infarct, even if those changes are not readily apparent to the naked eye. Furthermore, the use of attention mechanisms in some of these architectures assists with focusing the model’s attention on the most relevant parts of the image, further improving its accuracy. We will compare the performance of these four ML models in identifying infarct regions from LV strain data. The four ML models (Fig. 3) are described below in detail. Further specifics, including operation type, resolution size, and channel count, are provided in Table 1. After identifying the ML model with the highest performance among the four single-fidelity trained models, its performance was subsequently tested using human CMR data, which underwent a series of preparatory steps delineated in the <u>“Preprocessing of ML inputs”</u> section to predict infarct region of human patients. Following this test, further training was implemented by combining RCCM strains with human CMR strains in a multi-fidelity approach, utilizing the correlation between data obtained at varying levels of fidelity^47^ for improved performance of the ML model on human data.

**Table 1.** Detailed architecture of UNet, emphasizing the contracting and expansive paths, including block types, output resolutions, and channel numbers at different stages. The final output layer generates a single-channel image with a resolution the same as the input image.

| Contracting path |  |  | Expansive path |  |  |
| --- | --- | --- | --- | --- | --- |
| Block type | Output resolution | Output channel number | Block type | Output resolution | Output channel number |
| Input | 128 x 128 | 3 | Output | 128 x 128 | 1 |
| 2 blocks<br>Conv 3x3 | 128 x 128 | 16 | Conv 1x1 | 128 x 128 | 1 |
| Max pooling |  |  | 2 blocks<br>Conv 3x3 | 128 x 128 | 16 |
| 2 blocks<br>Conv 3x3 | 64 x 64 | 32 | Up-conv<br>Concatenate | 128 x 128 | 16 |
| Max pooling |  |  | 2 blocks<br>Conv 3x3 | 64 x 64 | 32 |
| 2 blocks<br>Conv 3x3 | 32 x 32 | 64 | Up-conv<br>Concatenate | 64 x 64 | 32 |
| Max pooling |  |  | 2 blocks<br>Conv 3x3 | 32 x 32 | 64 |
| 2 blocks<br>Conv 3x3 | 16 x 16 | 128 | Up-Conv<br>Concatenate | 32 x 32 | 64 |
| Max pooling |  |  | 2 blocks<br>Conv 3x3 | 16 x 16 | 128 |
| 2 blocks<br>Conv 3x3 | 8 x 8 | 256 | Up-Conv<br>Concatenate | 16 x 16 | 128 |

**Fig. 3.**
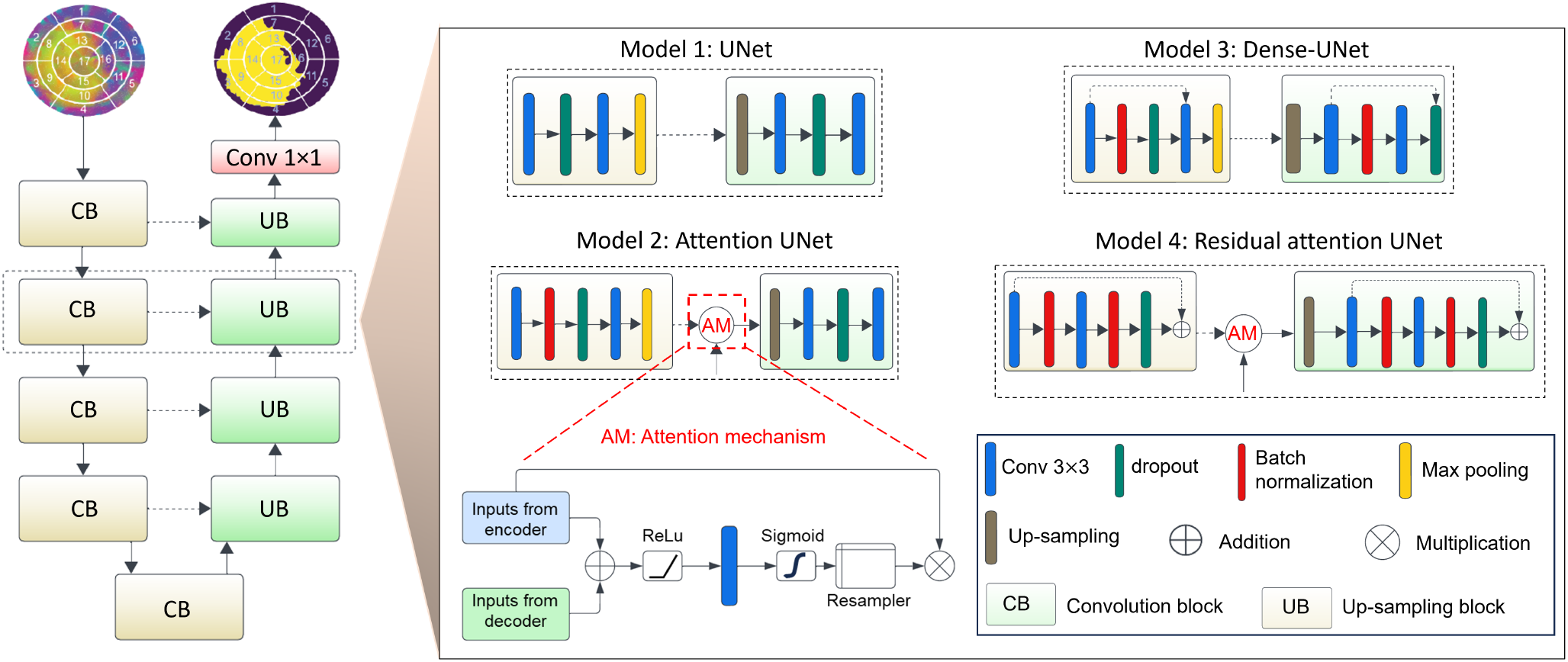
Illustration of base UNet and its variants for infarct segmentation in LV strain images. The base UNet architecture features a typical encoder-decoder structure. Attention UNet incorporates an attention mechanism in the encoder layers to focus on relevant features. Dense UNet adopts dense connections in the encoder to facilitate information flow. Residual attention UNet combines both residual connections and attention mechanisms for improved feature extraction and information preservation.

#### 2.3.1 UNet

UNet consists of symmetric contracting (encoder) and expansive (decoder) paths^33^. The contracting path comprises a series of convolutional layers, each followed by a rectified linear unit activation function and max-pooling layers. This arrangement progressively reduces the spatial dimensions of the feature maps while increasing the number of feature channels. This process helps the network to learn hierarchical representations of the input features, allowing it to extract relevant patterns and features. Conversely, the expansive path employs transposed convolutions to progressively upsample the feature maps, recovering spatial information and reconstructing the original input dimensions. Skip connections are employed between the encoder and decoder layers, enabling the fusion of high and low-resolution features from the encoder and decoder paths, respectively. These skip connections facilitate the precise localization of infarct regions by preserving fine-grained spatial details. The final layer of the UNet consists of a 1×1 convolutional layer followed by a sigmoid activation function. With a single-channel output, the convolutional layer condenses the multi-channel feature maps into a binary representation, where each pixel is assigned a value of either 0 or 1. The final layer predicts the probabilities of each pixel belonging to the infarct or normal region, with values closer to 1 indicating a higher likelihood of being part of the infarct and vice versa.

#### 2.3.2 Attention UNet

Expanding upon the foundation of UNet, the integration of an attention mechanism with the UNet architecture^48^ aims to unveil intricate, long-range dependencies crucial for enhancing infarct detection precision. While UNet effectively captures local features and spatial relationships, its reliance on convolutional operations alone may obscure subtle, interconnected patterns within the strain data. The attention mechanism serves as a valuable augmentation, introducing an additional module to scrutinize the feature map at each level. This module learns to assign importance scores to various spatial locations within the image, illuminating previously unseen regions of infarct. These importance scores guide the network to focus on the most informative strain patterns for infarct prediction. However, the advantages of attention are accompanied by computational overhead, potentially increasing model training time. Attention gates (AGs) are incorporated into the standard UNet architecture through skip connections, in which a gating vector from the decoder path is used for each pixel to determine focus regions. This gating vector includes contextual information for selectively reducing lower-level feature responses, a concept proposed in a prior study^49^, which utilizes AGs for the classification of natural images. We adopted additive attention^50^ to derive the gating coefficient, motivated by its higher experimental accuracy compared to multiplicative attention^51^, despite its computational cost.

#### 2.3.3 Dense UNet

Dense UNet incorporates dense connections, inspired by the DenseNet architecture^52^, to capture a more comprehensive picture of the underlying patterns. By forming a web-like structure of layers, Dense UNet creates an interactive environment where features interact seamlessly, refining one another and extracting more meaningful information^53^. Within each dense connection layer, a 3x3 convolution is followed by batch normalization and a ReLU activation. Feature maps from preceding layers are concatenated before input into the 3x3 convolution, fostering a comprehensive information integration. Similar to the standard UNet, Dense UNet incorporates skip connections that link the encoder and decoder paths, directly copying feature maps from the contracting path to the expanding path. This inclusion of dense connections may empower the model to comprehend features effectively, facilitating the capture of even fragmented or diffuse infarcts.

#### 2.3.4 Residual attention UNet

Lastly, residual attention UNet combines the strengths of two powerful techniques: residual connections and attention mechanisms for accurate infarct delineation^54^. Residual connections bypass the main data flow in the network, directly feeding the output of a previous layer to a subsequent layer. This helps to alleviate the vanishing gradient problem, leading to more stable training for deeper networks^55^. Additionally, the attention mechanism is incorporated to analyze feature maps at specific levels to guide the network’s focus toward relevant information. However, utilizing the residual attention UNet carries the risk of potential overfitting, posing a challenge to the model’s ability to generalize effectively to unseen data. This issue is of utmost importance in our specific context, as the unseen data specifically involves images from human CMR. This concern is particularly crucial in our context, where the unseen data pertains to human CMR imaging.

#### 2.3.5 Multi-fidelity deep neural network

Multi-fidelity deep neural network (DNN) based learning involves leveraging the relationship between data obtained at different fidelity levels^47^. In our study, our focus lies in capturing both linear and non-linear correlations between strains derived from low-fidelity RCCM and high-fidelity human CMR-LGE sources. Hence, we consider the generalized regressive scheme^34^ which is expressed as,

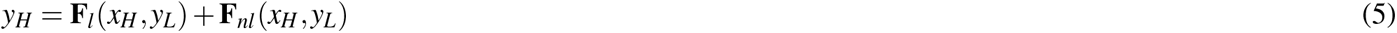

Here linear and nonlinear functions represented by **F**_*l*_ and **F**_*nl*_, respectively map the low-fidelity data to the high-fidelity data, learned by the neural networks, *NN*_*HF*1_ and *NN*_*HF*2_. Here, *y*_*L*_ is low fidelity output (infarct mask), *x*_*H*_ denotes high fidelity inputs (strains) and *y*_*H*_ is high fidelity output (infarct mask). The network architecture of the multi-fidelity deep neural network is shown in Fig. 4 composing of three neural networks. The first one *NN*_*LF*_ (*x*_*L*_, *θ*) is U-Net which is the same as previously discussed UNet in section 2.3.1, while the second NN *NN*_*HF*1_(*x*_*H*_, *y*_*L*_, *γ*_1_) is CNN kernel with 10×10 filter and third NN *NN*_*HF*2_(*x*_*H*_, *y*_*L*_, *γ*_2_) is also CNN kernel of same size but with leaky relu activation. Here, *θ, γ*_1_, *γ*_2_ indicates weights of *NN*_*LF*_, *NN*_*HF*1_ and *NN*_*HF*2_, respectively.In this study, to ensure a fair evaluation of the multi-fidelity deep neural network’s performance, we employed a fair training and testing protocol. Specifically, we utilized strains obtained from one human LGE-CMR (patient 1) with data augmentation (rotated to 90, 180, and 270 degrees) as high-fidelity training data as described in detail in the <u>“Preprocessing of ML inputs”</u> section, while reserving another set of strains (patient 2) for inference, thus preventing any data leakage between training and testing phases. We then reversed the roles of the samples, training the network with augmented data from patient 2 and using patient 1 data for inference, ensuring that the network’s performance was assessed on unseen data each time.

**Fig. 4.**
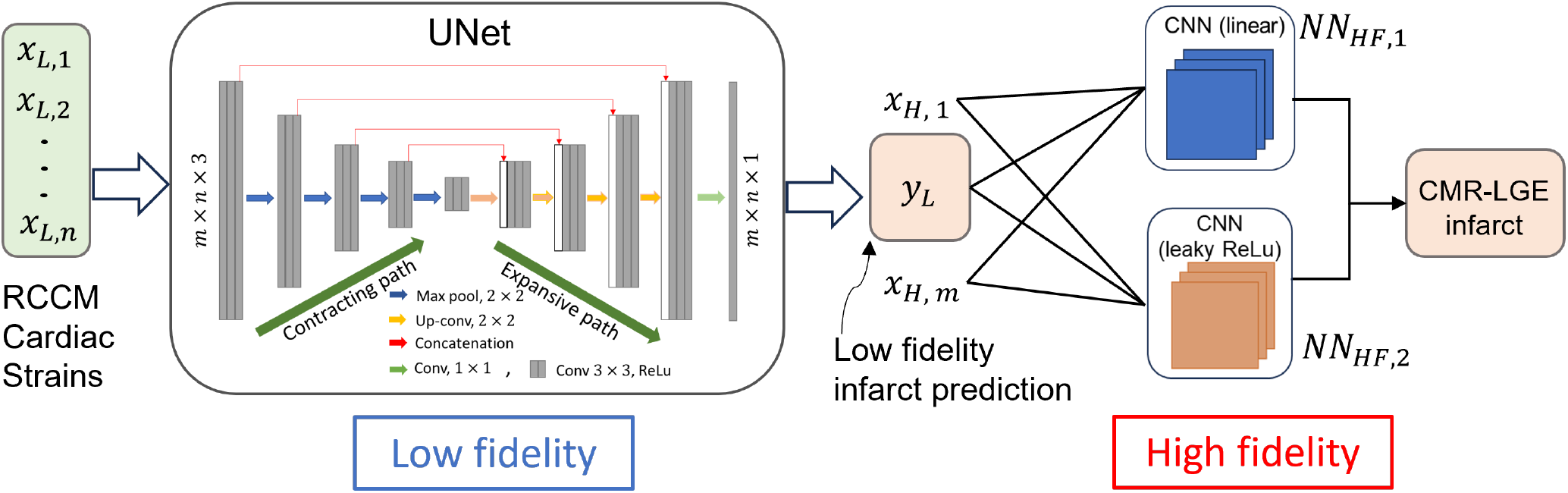
Network architecture highlighting the composite neural network that learns from the multi-fidelity data. The first block (UNet) constitutes the low-fidelity deep neural network *NN*_*LF*_ (*x*_*L*_, *θ*), followed by two high-fidelity models including *NN*_*HF*1_(*x*_*H*_, *y*_*L*_, *γ*_1_) and *NN*_*HF*2_(*x*_*H*_, *y*_*L*_, *γ*_2_), consisting of single CNN kernel without and with leaky ReLU activation functions, respectively. The combined output of the two high-fidelity neural networks is used to predict the binary infarct mask *y*_*H*_.

**Fig. 5.**
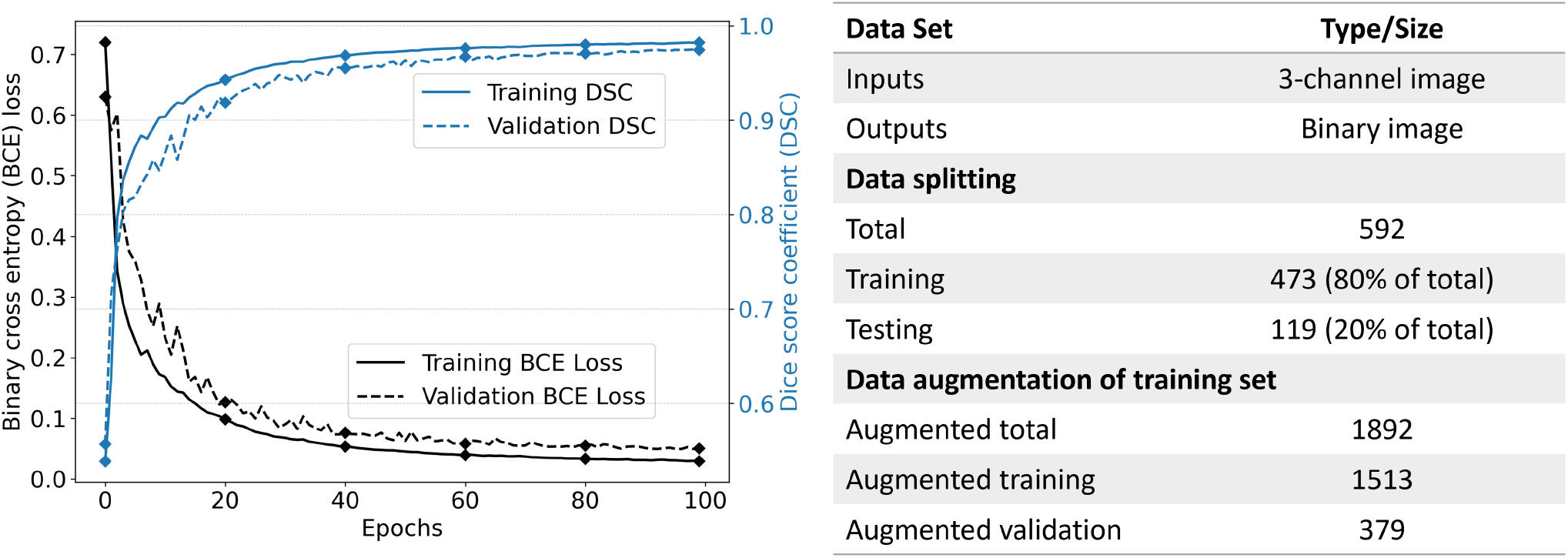
Training machine learning model using different loss functions including binary cross entropy (BCE), dice similarity coefficient (DSC) and the intersection over union (IoU) loss functions. 100 epochs, patience is 5, batch size is 128, image size is 128 by 128, and all images were normalized before training.

### 2.4 ML training process and optimization

As the attention UNet, dense UNet, and residual attention UNet were drawn from the UNet, we utilized the UNet ML model to explore general hyperparameters for identifying infarct regions. This exploration included carefully selecting input image dimensions, specifically 512×512, 256×256, and 128×128. A variety of loss functions, including binary cross-entropy (BCE) loss, dice similarity coefficient (DSC) loss, and the intersection over union (IoU) loss, were investigated to effectively minimize the dissimilarity between the predicted infarct regions and the corresponding ground-truth mask during training. The model performance evaluation was based on accuracy, precision, recall, IoU, and DSC evaluation metrics. The model was optimized using a gradient-based optimization algorithm (Adam) to minimize the chosen loss function. The following briefly describes the loss functions and evaluation metrics used in this work.

#### 2.4.1 Loss functions

The BCE loss is a commonly used loss function for binary pixel-wise segmentation tasks^56^. It measures the pixel-wise dissimilarity between the predicted segmentation map (*p*) and the corresponding ground-truth mask (*y*). The BCE loss is defined as:

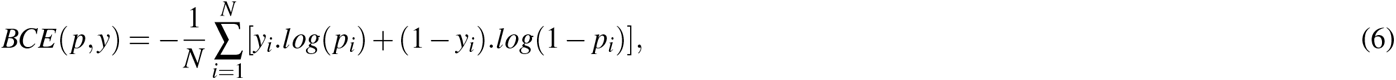

where *N* represents the total number of pixels in the segmentation map. The BCE loss penalizes incorrect predictions while encouraging the model to produce probabilistic values close to 1 for pixels in infarct regions and close to 0 for pixels in healthy regions.

The DSC loss is an additional loss function to enhance spatial overlap between the two masks. The DSC loss is defined as:

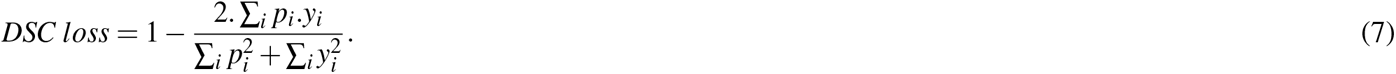

Similar to the DSC loss function, the intersection over union (IoU) is another commonly used loss function for segmentation. It also measures the spatial overlap between the predicted segmentation map (*p*) and the ground-truth mask (*y*). The IoU loss is given by:

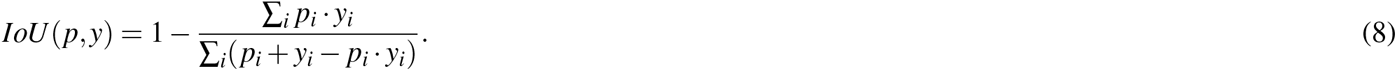

By optimizing the DSC and IoU loss functions, the model learns to produce precise segmentation maps that accurately delineate infarct regions. In the above equations, *p*_*i*_ represents the pixel value in the predicted map at *i*th pixel, and *y*_*i*_ is the pixel value in the ground-truth mask at *i*th pixel. The summation spans all pixels in the image. A smaller loss function value indicates improved segmentation accuracy, signifying greater overlap between the predicted and true masks.

The trainable variables for the multi-fidelity neural network were learned by minimizing the following loss function which was minimize using the Adam optimizer:

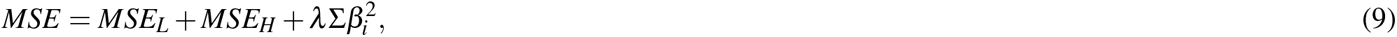

where

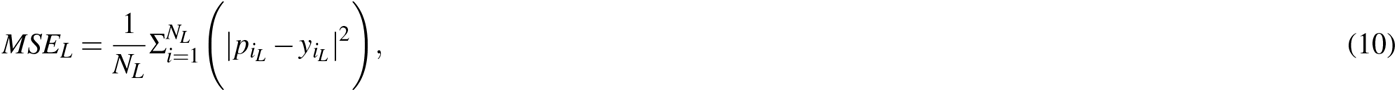

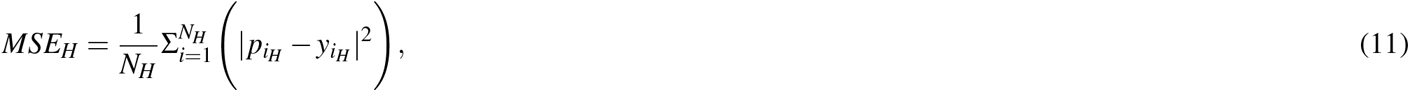

Here, *N*_*L*_ indicates the total number of training samples in low-fidelity dataset, and *N*_*H*_ indicates the total number of training samples in the high-fidelity dataset. 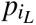 and 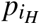 denotes the predicted outputs of the *NN*_*L*_ and *NN*_*H*_, *β* is any weight in *NN*_*L*_ and *NN*_*H*_ and *λ* is the L2 regularization rate for *β*.

#### 2.4.2 Evaluation metrics

Several well-known metrics, including accuracy, precision, and recall, were used to evaluate models’ performance to segment infarct regions, which provided important insights into the models’ overall effectiveness for prediction. However, two additional metrics, namely IoU and DSC, hold particular significance for infarct region segmentation tasks, where the spatial overlap is critical. These metrics, described below, are crucial for assessing how well the model can locate the infarct regions within the LV myocardium.

The IoU, also called the Jaccard index, quantifies the extent of the region where the ground-truth mask and predicted segmentation map overlap. It is determined by dividing the intersection of the predicted and actual masks by their union:

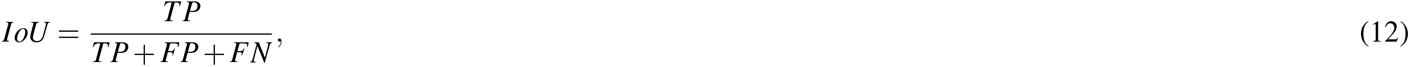

where TP represents true positive (correctly predicted infarct pixels), FP is false positive (incorrectly predicted infarct pixels), and FN is false negative (infarct pixels missed by the model). The predicted and ground-truth masks must perfectly overlap for the IoU to be 1, which ranges from 0 to 1. Increased segmentation accuracy is shown by a higher IoU score, which measures the degree of spatial overlap between the model’s predictions and the actual infarct regions.

The DSC, also referred to as the F1-score, is another metric that measures the spatial overlap between the predicted infarct region with the ground-truth infarct region. It is defined as twice the intersection of the predicted and ground-truth masks divided by their sum:

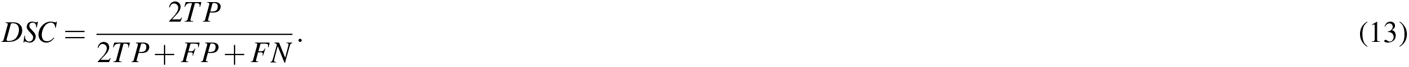

Similar to the IoU, the DSC ranges from 0 to 1, with 1 representing a perfect segmentation. DSC is particularly useful in evaluating the segmentation accuracy of smaller infarct regions, as it considers the relative contribution of true positive, false positive, and false negative predictions.

### 2.5 Human LGE-CMR used for validation

Human LGE-CMR data was collected from two patients (n=2) with confirmed MI. All personal identifiers were removed in accordance with the National Institute of Health de-identification protocol, comprising 18 elements considered to be protected health information (https://privacyruleandresearch.nih.gov/pr_0_8.*asp*). The research protocol was approved by the Houston Methodist Research Institute review board. Briefly, cine CMR scans were acquired using a 3.0-T clinical scanner (Siemens Verio; Siemens, Erlangen, Germany) with phased-array coil systems. CMR scans included a short axis stack using a steady-state free-precession sequence. LGE images were acquired over slice positions matched to cines about 10 to 15 minutes after intravenous gadolinium-based contrast administration (gadopentetate dimeglumine, gadoterate meglumine; 0.15 mmol/kg) with in-plane spatial resolutions of 1.8 mm by 1.3 mm and slice thicknesses of 6 to 7 mm with 3 to 4 mm gap. Additional details of LGE-CMR scans are available in^57,58^. ML models trained on only RCCM data (single-fidelity) were evaluated by predicting infarct regions of both human patients. However, for a multi-fidelity approach, strains from one LGE-CMR scan were used in training, and another was used for evaluation due to a scarcity of human LGE-CMR scans. This procedure was repeated for both human patients alternately. The strains calculated from CMR scans and preprocessed, as described in <u>“Preprocessing of ML inputs”</u> section, were used as inputs to the models to predict MI regions. The CRL strains from CMR slices were calculated at ES, using the end-diastolic frame as a reference point. To achieve this, an optimization-based image registration approach^59–61^ was implemented. The registration determines the optimal transformation that aligns a reference frame, denoted by *ℱ*, with a moving image, denoted by *ℳ*. This alignment was achieved by minimizing the cost function, denoted by *W*, as expressed through the following equation:

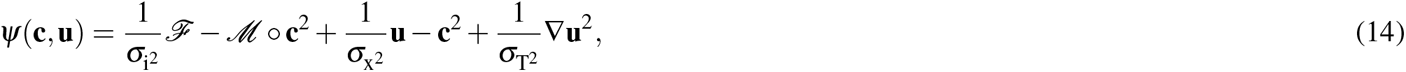

where *σ*_i_ and *σ*_x_ represent the noise intensities, while *σ*_T_ serves as a regularization factor. The term **u** denotes the parametric transformation, representing the Cartesian displacement vector at each pixel between two consecutive time frames. Simultaneously, **c** signifies the corresponding non-parametric spatial transformation.

The large deformation theory was employed to derive the Green-Lagrange strain tensor (**E**) for Cartesian displacements obtained from imaging as pointwise data. The strains were computed utilizing the total deformation gradient (**F**) and the identity matrix (**I**) in accordance with the expression: 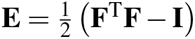. Here, **F** was determined as the propagation of the deformation gradient between two consecutive load increments (**F**_i_), given by:

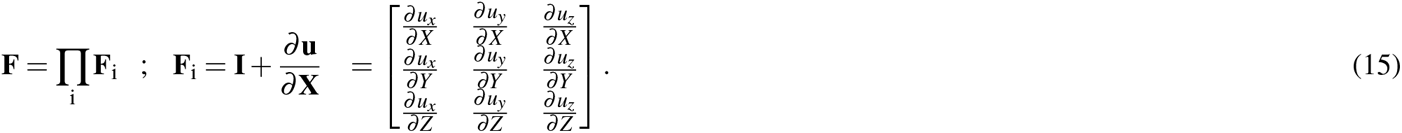

Here, u_x_, u_y_, and u_z_ denote the displacements in the x, y, and z directions between two consecutive load increments. Finally, CRL strains were obtained by transforming the Cartesian strains into CRL axes using an orthonormal transformation matrix (**Q**) as:

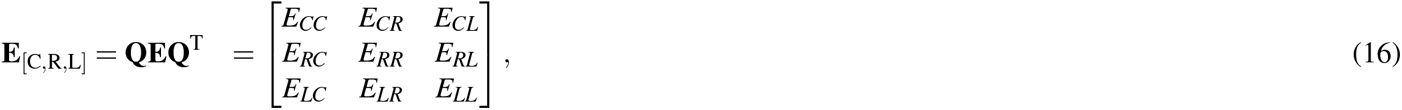

where *E*_*CC*_, *E*_*RR*_, and *E*_*LL*_ represent the circumferential, radial, and longitudinal strains, respectively.

## 3 Results

A comprehensive analysis of the predictive capability of the encoder-decoder based ML models for infarct region identification from LV strain was conducted. We began with an extensive examination of different image sizes in the ML model to determine the optimal image size for training to enhance the ability of the model to capture fine-grained features. This comparison was carried out under the same experimental conditions, considering the uniform data preprocessing and utilization of the same loss function. Next, we investigated the impact of various loss functions on the performance of the ML model while keeping the image size fixed. These loss functions, including the BCE, DSC, and IoU loss, were evaluated using carefully chosen evaluation metrics. Furthermore, the efficacy of all chosen encoder-decoder-based architectures were compared against each other. We selected the best performing architecture among all as single-fidelity DNN to further extend the framework to multi-fidelity DNN. Finally, we tested the multi-fidelity trained DNN in predicting the infarct region from human LGE-CMR-derived strains. This evaluation aimed to demonstrate the model’s effectiveness when applied to data from human patients, thereby evaluating its capacity for generalization and its suitability for clinical application.

### 3.1 Influence of image size on model adaptability

We considered three distinct image sizes for experimentation: 128×128, 256×256, and 512×512 pixels. Each image size was preprocessed using the steps detailed in the Methods section, and the BCE loss function was employed consistently to ensure fair and unbiased comparisons. The results of this investigation are presented in Table 2(a), where the performance metrics, including accuracy, precision, recall, IoU, and DSC scores, are listed for each image size. Our analysis revealed a remarkable consistency in model performance across varying image sizes. The IoU and DSC metrics exhibited minimal differences, underscoring the model’s adaptability across image sizes. Notably, even the 128×128 image size, which is often associated with reduced spatial information, provided IoU (0.9757) and DSC (0.9877) values indicative of effective infarct region localization. The 256×256 and 512×512 image sizes also showcased comparable spatial overlap and segmentation accuracy results. Given this consistency in performance, it was deemed justifiable to make use of the 128×128 picture size for later training stages. Using a reduced image size reduced computation and training time while maintaining high accuracy.

**Table 2.** Comparison of different image sizes and loss functions in UNet ML models.

| (a) Different image sizes in UNet |  |  |  |  |  |
| --- | --- | --- | --- | --- | --- |
| Image sizes | Accuracy | Precision | Recall | IoU | DSC |
| 128×128 | 0.9874 | 0.9925 | 0.9829 | 0.9757 | 0.9877 |
| 256×256 | 0.9884 | 0.9950 | 0.9801 | 0.9753 | 0.9875 |
| 512×512 | 0.9882 | 0.9937 | 0.9798 | 0.9738 | 0.9867 |

| (b) Different loss functions in UNet |  |  |  |  |  |
| --- | --- | --- | --- | --- | --- |
| Loss functions | Accuracy | Precision | Recall | IoU | DSC |
| BCE loss | 0.9835 | 0.9941 | 0.9737 | 0.9681 | 0.9837 |
| IoU loss | 0.9815 | 0.9785 | 0.9856 | 0.9648 | 0.9821 |
| DSC loss | 0.9843 | 0.9844 | 0.9849 | 0.9698 | 0.9847 |

### 3.2 Quantitative analysis of loss functions

The effectiveness of several loss functions, namely BCE, DSC, and IoU loss, was examined in UNet. The quantitative analysis was performed by comparison of metric scores, including accuracy, precision, recall, IoU, and DSC for each loss function (Table 2 (b)). All loss functions exhibited comparable accuracy. Specifically, BCE obtained a DSC of 0.9837 and an IoU of 0.968, whereas the DSC loss function displayed a DSC of 0.9847 and an IoU of 0.9698. The IoU loss function produced a DSC of 0.9821 and an IoU of 0.9648. For brevity, all ML predictions of the FE results shown in Figs. 6 and 7 were based on the training using the BCE loss function. However, models trained with different loss functions were subsequently evaluated on human CMR-derived strains (Section 3.4).

**Fig. 6.**
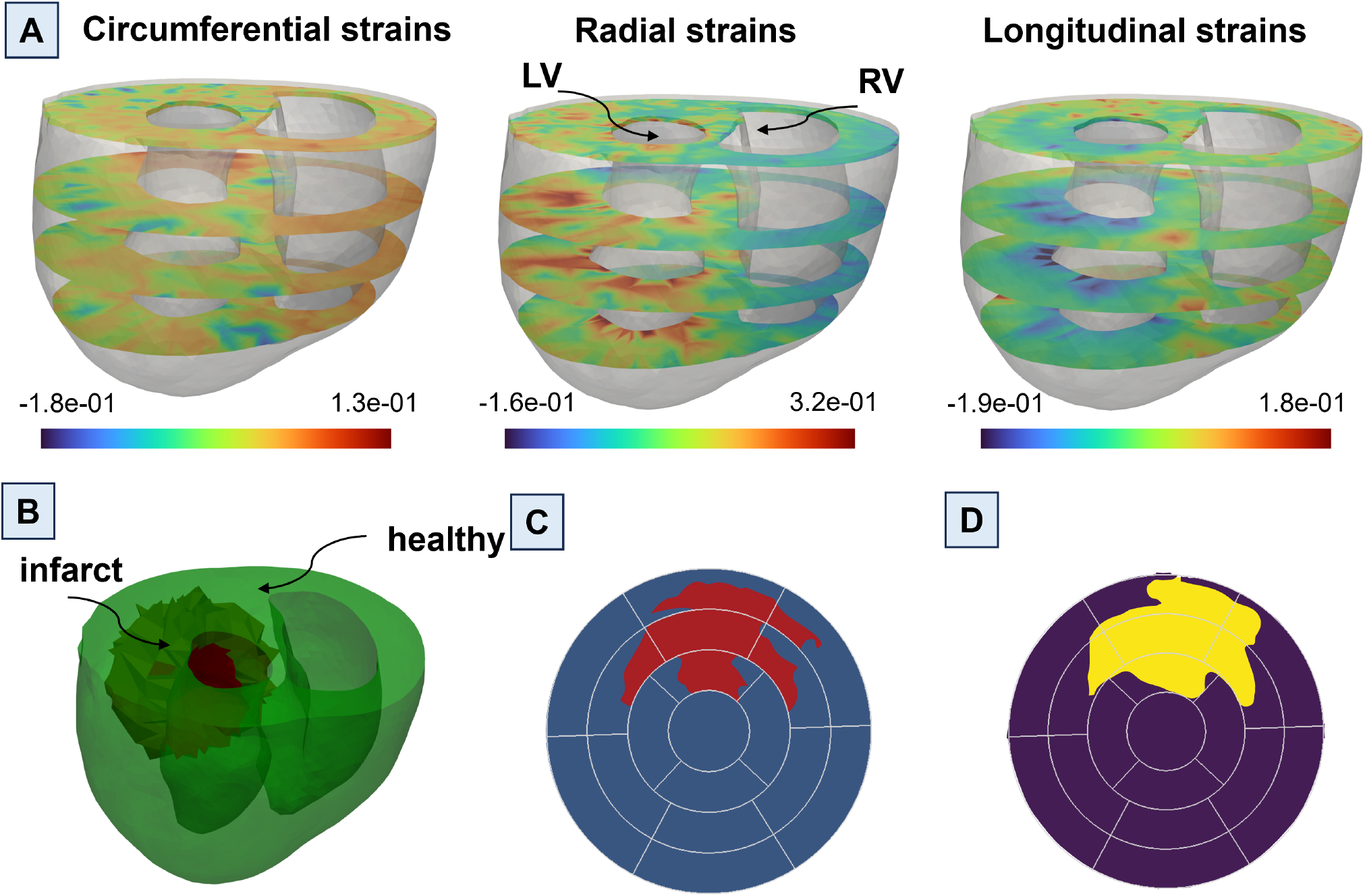
A representative example showcasing the prediction of the infarct region using the UNet machine learning (ML) model based on cardiac circumferential, radial, and longitudinal (CRL) strains obtained from finite element (FE) simulation. (A) Cardiac CRL strains depicted at the base, mid, apical, and apex of the left ventricle. (B) A 3D image of the heart model, where healthy regions are represented in green and the infarct region in red. (C) An American Heart Association (AHA) bullseye map derived from the 3D heart model representation, indicating healthy (blue) and infarct (red) regions. (D) ML-predicted healthy (purple) and infarct (yellow) in the AHA bullseye map format.

**Fig. 7.**
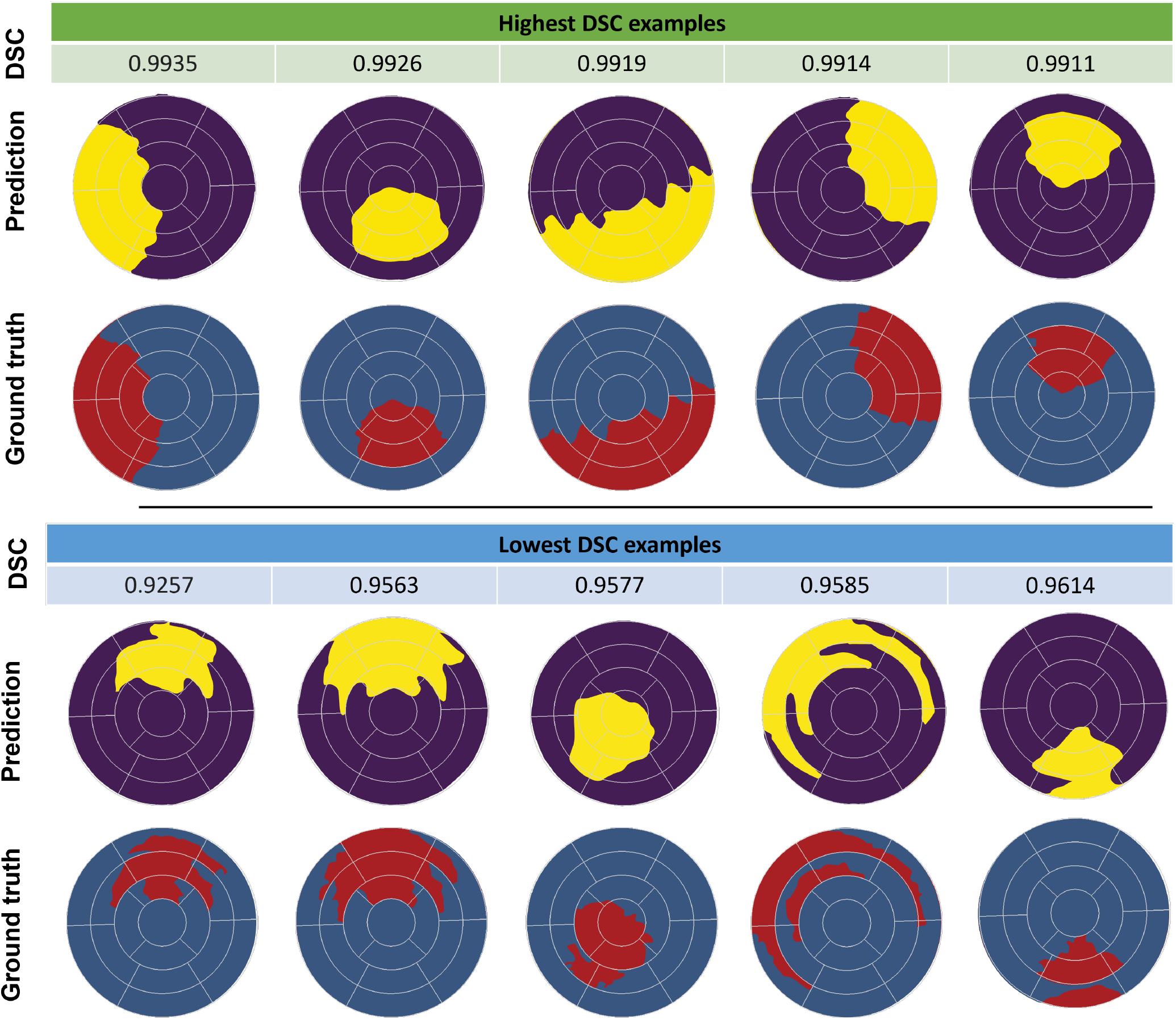
Visual comparison of single-fidelity finite element (FE) ground truth against UNet predictions, showcasing the highest and lowest dice similarity coefficient (DSC) scores. Five examples each highlight the model’s exceptional accuracy in infarct region delineation, revealing both optimal and challenging predictions.

### 3.3 Impact of different architectures on performance metrics

All four distinct models, namely UNet, attention UNet, dense UNet, and residual attention UNet, required trainable parameters in the order of millions (Table 3). UNet demonstrated a notable reduction in parameter count compared to other models, while all utilized models remained parameter-efficient compared to other segmentation models^62–64^. Although UNet streamlined its parameter count, its performance in infarct region identification remained exceptional. Evaluation metrics in Table 4 supported this observation, with UNet achieving an IoU of 0.968 and DSC of 0.9837. Similarly, attention UNet, dense UNet, and residual attention UNet showed comparable metrics to UNet, although with slight variation in performance attributed to their distinct architectures. These results highlighted the balance between computational efficiency and segmentation accuracy in the UNet model. A representative example of the input data, ground truth infarct region, and UNet model prediction for one heart model is illustrated in Fig. 6. Additionally, Fig. 7 showcases a comparison between ground truth and prediction for examples with the highest and lowest DSC metric scores, demonstrating minimal variation between the predicted infarct regions and the ground truth masks even in examples with the lowest DSC scores. These findings emphasize the effectiveness of our approach, highlighting that the computationally less intensive UNet can achieve comparable high metrics when compared to attention UNet, dense UNet, and residual attention UNet architectures.

**Table 3.** Comparative analysis of trainable parameters in encoder-decoder-based ML architecture, illustrating the model complexity in relation to the increasing number of parameters.

| ML models | Number of trainable parameters |
| --- | --- |
| UNet | 1.94 million |
| Attention UNet | 2.38 million |
| Dense UNet | 2.70 million |
| Residual attention UNet | 3.71 million |

**Table 4.** Comprehensive evaluation of all the machine learning models across different metrics.

| ML Model | Accuracy | Precision | Recall | IoU | DSC |
| --- | --- | --- | --- | --- | --- |
| UNet | 0.9835 | 0.9940 | 0.9737 | 0.9680 | 0.9837 |
| Attention UNet | 0.9839 | 0.993 | 0.9756 | 0.9689 | 0.9842 |
| Dense UNet | 0.9893 | 0.9939 | 0.9851 | 0.9792 | 0.9895 |
| Residual attention UNet | 0.9876 | 0.9935 | 0.9823 | 0.976 | 0.9879 |

### 3.4 ML model performance assessment with human LGE-CMR

To rigorously evaluate our ML model performance and indicate the feasibility of the presented approach in the clinical setting, we used cardiac strain data from human LGE-CMR scans that is accessible in clinical scenarios. By applying our trained ML model to the strain data calculated using image registration from CMR scans, we obtained predictions that were subsequently compared against the ground truth mask obtained from LGE-CMR scans (Fig. 8). This comparison enabled a comprehensive evaluation of our model predictive accuracy in identifying the precise infarct regions within the LV. The model performance was evaluated using the DSC metric score. The resulting DSC obtained from single-fidelity trained ML models were 0.755 ± 0.0715 and 0.762 ± 0.0482 for first and second patients, respectively (Fig. 8c). The DSC scores derived from multi-fidelity trained ML models were notably higher, measuring 0.873 ± 0.0099 for the first patient and 0.819 ± 0.0145 for the second patient (Fig. 8d). This result highlights the effectiveness of the multi-fidelity approach in delineating infarct regions, attesting to the possibility of simulation-enhanced ML tools in the clinical setting for predicting the presence and location of infarcts within the LV using standard cine CMR scans.

**Fig. 8.**
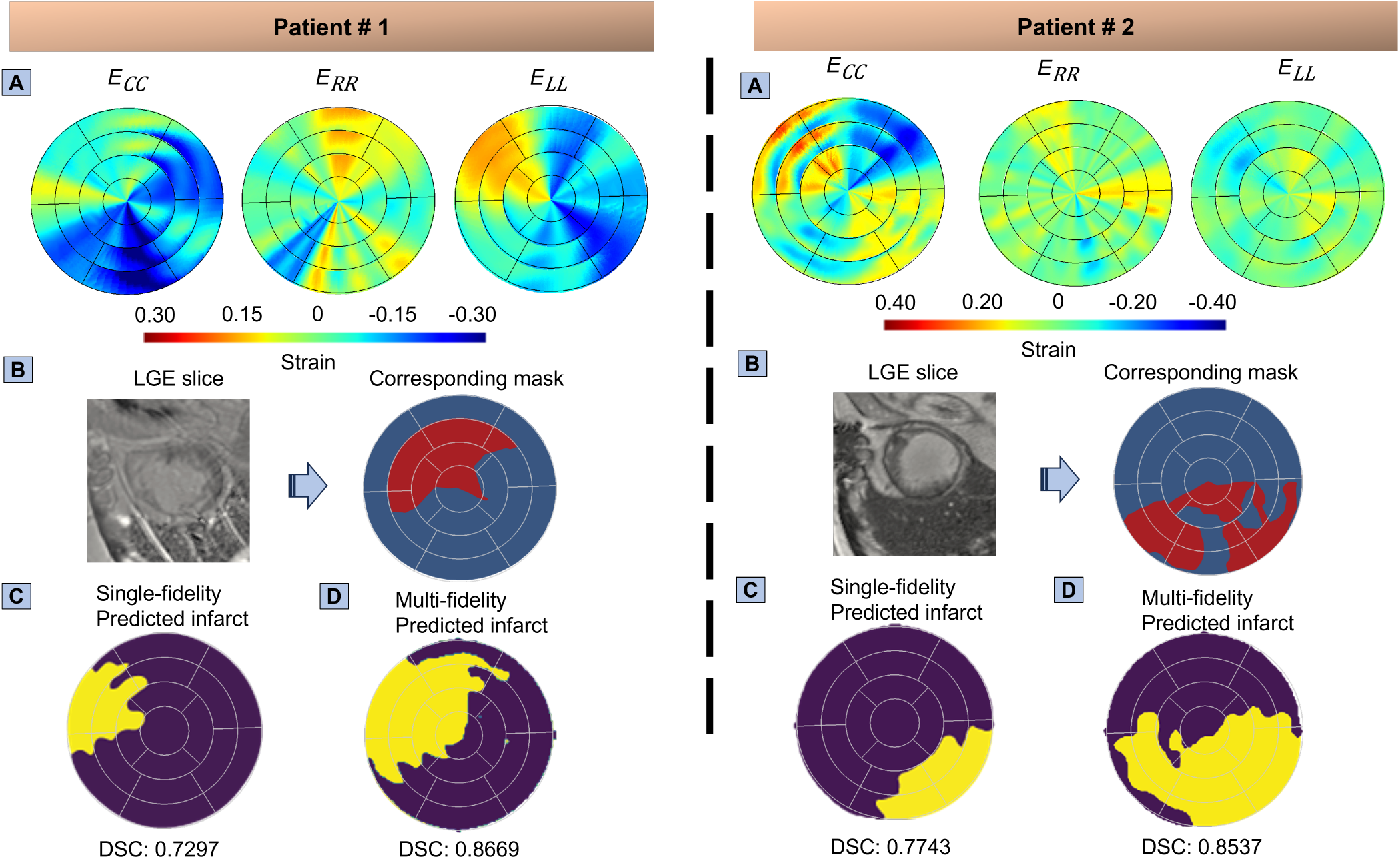
Human late gadolinium-enhanced cardiac magnetic resonance (LGE-CMR) imaging illustrates cardiac circumferential, radial, and longitudinal (CRL) strains alongside predictions of infarct regions generated by both single-fidelity and multi-fidelity machine learning (ML) models, in comparison with ground truth derived from gadolinium contrast agent. The components of this illustration includes (A) CRL strains extracted from LGE-CMR imaging, (B) One of the infarct LGE slices from the entire stack of images, accompanied by a representation of healthy and infarct regions on an American Heart Association (AHA) bullseye map, serving as ground truth derived from LGE-CMR, (C) Dice similarity coefficient (DSC) scores for the single-fidelity trained ML model applied to the first and second patients, yielding scores of 0.755 ± 0.0715 and 0.762 ± 0.0482 respectively. Predictions from a single ML run yielded DSC scores of 0.7297 and 0.7743 for the respective patients. (D) DSC scores for the multi-fidelity trained ML model applied to the first and second patients, resulting in scores of 0.873 ± 0.0099 and 0.8193 ± 0.0145 respectively. The ML predictions associated with these scores demonstrate DSC values of 0.8669 and 0.8537 respectively.

### 3.5 Impact of number of training samples in single-fidelity and multi-fidelity learning-based models

We consider a multi-fidelity architecture making use of four different values of low-fidelity (RCCM) data *N*_*LFR*_ = [20, 200, 400, 592] in training to assess the performance of a multi-fidelity DNN compared to a single-fidelity UNet. We evaluated the performance of both approaches with five random seed initialization for each value of *N*_*LFR*_ to compute the mean and standard deviation of DSC of human patients (Fig. 9). Our findings clearly highlighted that the multi-fidelity approach outperformed in terms of accuracy and has less model uncertainty in the overall prediction capability of the infarct mask, even with very limited high-fidelity data (in our case, just one sample of human LGE-CMR). This ablation study additionally aimed to demonstrate how low-fidelity (RCCM) data augmented the overall learning of the multi-fidelity DNN. We observed a saturation point in the DSC at *N*_*LFR*_ = 400, with a slight increase in DSC corresponding to *N*_*LFR*_ = 592. This suggests that beyond *N*_*LFR*_ = 400, there is no further correlation that the model can exploit from the low-fidelity data, even with an increased number of RCCM samples. Utilizing a multi-fidelity approach, we achieved DSC values of 0.873 ± 0.0099 and 0.8193 ± 0.0145 for patient 1 and patient 2 respectively, when trained with the highest number of low-fidelity samples (*N*_*LFR*_ = 592) considered in this study.

**Fig. 9.**
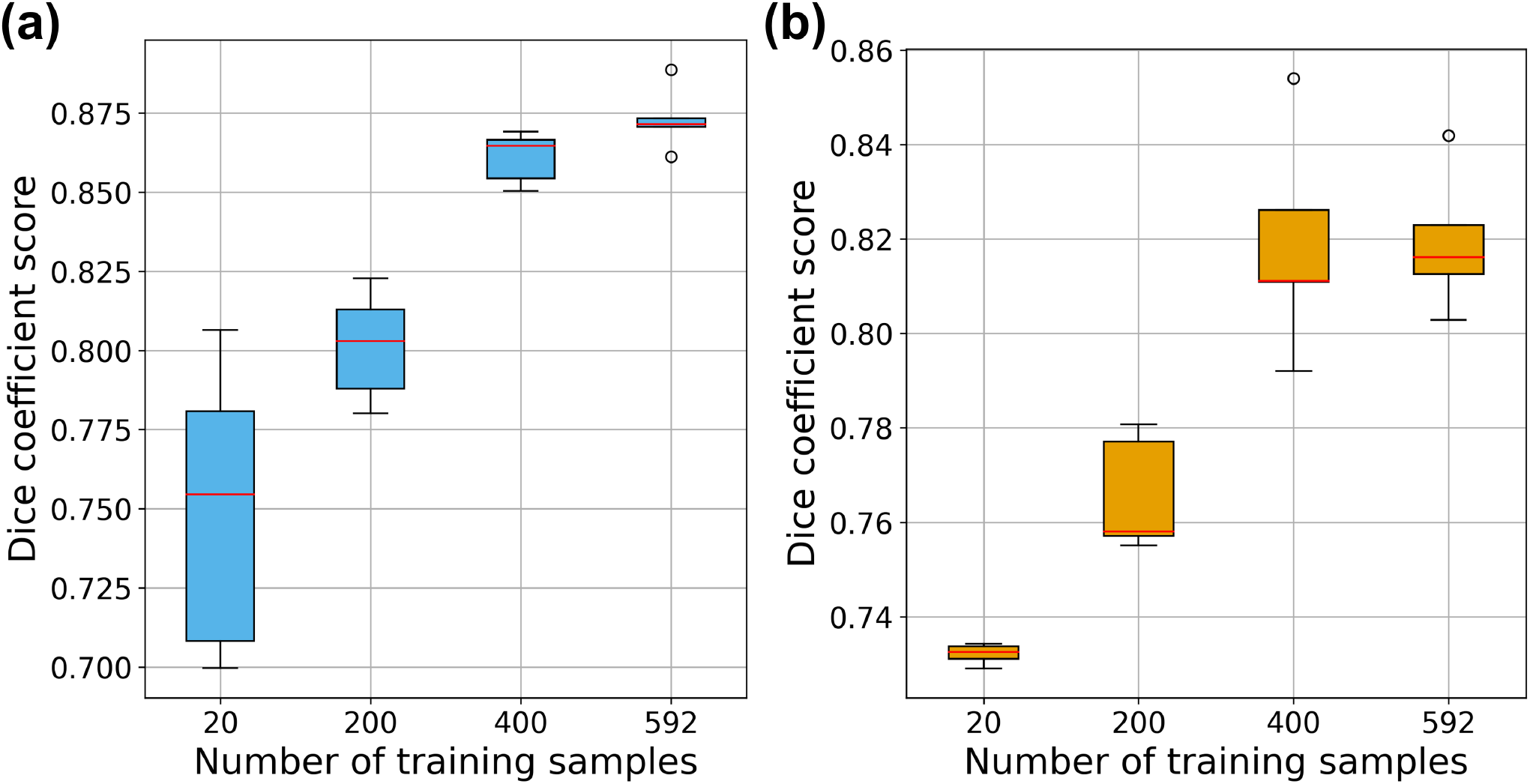
The impact of the number of low fidelity training samples (RCCM based) on the prediction of multi-fidelity learning based models: (a) first human patient, and (b) second human patient.

## 4 Discussion

### 4.1 Non-invasive identification of infarct regions in the LV

Our approach capitalized exclusively on image-derived cardiac strain data as input features for our ML model. Through seamlessly integrating cardiac strain images with ML techniques, we enabled the precise localization of infarct tissue by extracting both local and global image features. This innovative framework bypassed the need for the invasive administration of GCA, addressing a significant limitation in traditional methods. The avoidance of GCA injection not only mitigates potential risks and patient discomfort but also minimizes potential adverse effects associated with GCA. Our approach thus offers a promising avenue for accurate infarct identification without subjecting patients to invasive procedures or GCA-related concerns. This advancement promises to redefine non-invasive cardiology diagnostic strategies and optimize patient safety and clinical decision-making.

### 4.2 Harnessing image-based strain representations in convolution-based networks

The key query addressed in our research was the effective incorporation of cardiac strain data into ML models to identify infarct location. We solved this problem by converting strain data into an image-based format rather than using numerical strain values directly. This approach was employed to utilize the capabilities of convolutional-based networks, which are well-known for their ability to handle pixel-based data in tasks such as object categorization, object recognition, and scene interpretation. We harnessed the power of convolution-based networks to examine and grasp pixel values from the complex images of strain data. Furthermore, direct utilization of numerical values of strain data was avoided in the ML model due to inconsistencies in numerical datasets from diverse CMR cases caused by the inherent variability in pixel distribution among CMR images. More specifically, we used the AHA segmentation procedure as a first step before translating the data into standardized images. This dual transformation assured uniform image sizes and spatial arrangements, which not only regularized the input features with ML models but also allowed the seamless integration of data augmentation approaches, which was a key component of model generalization. In contrast to the numerical values with tree-based models or fully connected networks, convolution-based networks captured relevant features directly from raw pixel values. Furthermore, this approach eliminated the need for manual feature engineering, a critical advantage that ideally aligns with the transition to image-based strain representations.

### 4.3 Utilization of CRL strains for enhanced infarct region identification

Our approach incorporated cardiac CRL strains in the inputs of ML models with a limited number of channels using pre-processing steps to account for a wide range of possibilities of infarct regions. The trained model did not only show promise in the non-invasive identification of the infarct scar sites but may also help identify injured regions of the myocardium with reduced function, such as surrounding regions closely connected with existing infarct zones. Notably, this comprehensive use of strain data enabled the localization of active infarct locations and the characterization of “border” zones with compromised contractility compared to the LGE-CMR approach, which can primarily locate mature scar regions. This contrast was evident in Fig. 8 where GCA was not spread enough to locate the infarct region accurately, but our ML model still found the infarct region and assisted with estimating the extent of the border regions at risk of losing contractility, thus improving the prognosis of the transition to heart failure in MI patients. In essence, using processed cardiac strains as markers to locate infarct regions not only reduces the risk of invasive procedures by avoiding GCA injection but also promises to improve the stratification of MI at a higher risk of transitioning to systolic heart failure. Furthermore, our approach used cardiac strain data which can be obtained from more ubiquitous and affordable imaging modalities than CMR, such as echocardiography, reinforcing the possibility of implementation of such approach in clinical settings.

### 4.4 Pursuit of best ML model to use cardiac strains

Among the single-fidelity ML models investigated in this study, UNet exhibited comparable DSC with its variants in identifying infarct regions in human CMR strain data. However, the observed performance of UNet and its variants on human CMR is attributed to their limits in generalization across diverse datasets. All UNets demonstrated promising performance on FE simulation data (Table 4), however their challenge to effectively transition to human CMR data is noteworthy (Fig. 8). The lower accuracy of these single-fidelity frameworks on human LGE-CMR data highlights the difficulty of transitioning trained models from synthetic data. We attempted to overcome such challenge with a multi-fidelity ML approach. Our results indicate that for every *N*_*LFR*_ value, DSC for multi-fidelity ML approach outperformed single-fidelity based models (Fig. 9). The noted improvement after the addition of just one set of human data indicates the promise of developing accurate multi-fidelity simulation-based ML models with small amounts of human data, which is often difficult to obtain and process. All models, including multi-fidelity DNN, could additionally be extended to incorporate the physical laws governing the myocardial behavior^65^ which could potentially augment the accuracy and also help in reducing the number of training samples required in both low- and high-fidelity datasets. More sophisticated architecture of multi-fidelity DNN could be analyzed like residual multi-fidelity DNN^66^ with a slightly increased number of high-fidelity realizations. This presents a potential avenue for future research, aiming to refine the performance and adaptability of these models to efficiently and reliably address the sub-optimal strain quantification from human CMR imaging.

### 4.5 Detailed cardiac mechanics from 4D cardiac strains

Although our study focused on infarct region identification, it laid the groundwork for various possible future directions. While we exclusively provided ES strains on specific cardiac slices aligned with the AHA segmentation, it is possible to obtain complete spatiotemporal (four-dimensional or 4D) strains in the LV for training, giving our ML model richer inputs for capturing LV mechanics and locating infarct and compromised regions accurately. This expansion could constitute a significant improvement over the common utilization of cardiac strains in the form of a limited number of short-axis slices, keeping in mind that LGE-CMR acquisitions on short-axis slices could, at best, locate infarct locations at those specific slices, whereas complete spatial strain quantification can more precisely locate infarct in the LV. Furthermore, integrating strains obtained at all time points of the cardiac cycle will open new possibilities for extracting complex insights beyond locating infarct regions. This temporal dimension allows for understanding the difference between infarct versus infarct tissues not only in contractile behavior but also in relaxation and passive behaviors, assisting with determining their distinct mechanical characteristics. For instance, by analyzing the entire cycle, we may be able to distinguish between the contributions of collagen and myofibers to tissue stiffness, presenting an opportunity to improve the understanding of the passive behavior of infarct tissues. In essence, using 4D strains offers to improve the full characterization of myocardial scars and subsequent prognostic and therapeutic approaches in MI patients.

### 4.6 Limitations

Encouraged by the promise of using cardiac strains to identify infarct regions, addressing the four limitations listed below can further advance the translation of our tool to the clinic:

#### (i) Challenges with human LGE-CMR validation

Some inherent limitations of human LGE-CMR imaging may introduce a unique set of discrepancies between the predictions by our models and those by LGE. In particular, gadolinium enhancement may not present the complete infarct location since LGE-CMR acquisition captures infarct on selective short-axis slices and is best at capturing mature scar tissues. In situations when the LGE distribution does not sufficiently capture the extent of infarct tissue, our model’s prediction of the infarct region may differ from those by LGE-CMR presumed to be the ground truth. Therefore, future rodent studies consisting of in-vivo CMR and ex-vivo infarct identification using histology can serve as a critical benchmark to evaluate and further refine our model’s prediction.

#### (ii) Preprocessing for strain-based infarct identification

Although our approach accurately estimates infarct location, it also adds a crucial intermediary step to extract strains from segmented slices, which introduces complexity before providing to the ML model. In an alternative scenario, one could envision directly utilizing the individual slices at all time points from end-diastole to ES as inputs for our ML model. The model could potentially infer and localize infarct regions by leveraging pixel movements across time frames. While this alternative path might offer a seemingly streamlined process, it will raise a serious issue about the probable loss of subtle details necessary for precise infarct identification. In the direct utilization of imaging slices, omitting explicit strain extraction, the model may unintentionally neglect details of heart mechanics and tissue behavior that are adequately captured through strain analysis if it only considers pixel motions.

#### (iii) Comprehensive three-dimensional (3D) infarct localization

Our methodology was developed to be consistent with the cardiac imaging modalities that provide data in the form of 2D slices. Although our approach can identify infarct regions on those slices, there will be a need for an additional but straightforward extension of the method to map the full 3D extent of the infarct region, which will provide a more reliable prognostic marker than 2D representations of infarct regions. Indeed, the prediction of infarct region from cardiac strains, inherent in our method, perfectly lends itself to the possibility of 3D reconstruction of infarct regions as long as rigorous strain calculation algorithms can determine complete spatiotemporal strains through proper interpolation of cardiac motion across 2D slices^60,61^.

#### (iv) Integration of machine learning with multi-scale modelling

In our study, we successfully demonstrated the ability of ML-based frameworks to identify the infarct region from CMR strains. However, relying solely on ML overlooks essential physical laws, leading to potentially ill-defined problems or solutions that defy physics. Multi-scale modeling, a well-established approach, combines data from various scales and physical phenomena to reveal the underlying mechanisms of function. In the past two decades, this approach has proven effective in constructing detailed organ models by integrating knowledge across tissue, cellular, and molecular levels. The recent advancements in ML, with its capacity to handle diverse, multi-fidelity data and uncover complex correlations, offer a unique opportunity in this context. Nevertheless, multi-scale modeling on its own often struggles to efficiently merge vast datasets from different sources and resolutions. By combining ML with multi-scale modeling, we can create robust predictive models that incorporate fundamental physics, address ill-defined problems, and navigate extensive design spaces^67^. Adopting a multidisciplinary approach that blends ML and multi-scale modeling can unlock new insights into disease mechanisms, aid in identifying novel therapeutic targets and strategies, and support decision-making processes for improving human health.

## Acknowledgements

This work was supported by the National Institutes of Health awards R00HL138288 to R.A. and R01HL168473 to N.K. and G.K. and the National Science Foundation award 2244995 to R.A.

## Data availability

The data that support the findings of this study are available from the corresponding author, R.A., upon request.

