## Supplementary Fig. 1 for "Multi-Modality Deep Infarct: Non-invasive identification of infarcted myocardium using composite in-silico-human data learning"

### Supplementary Information

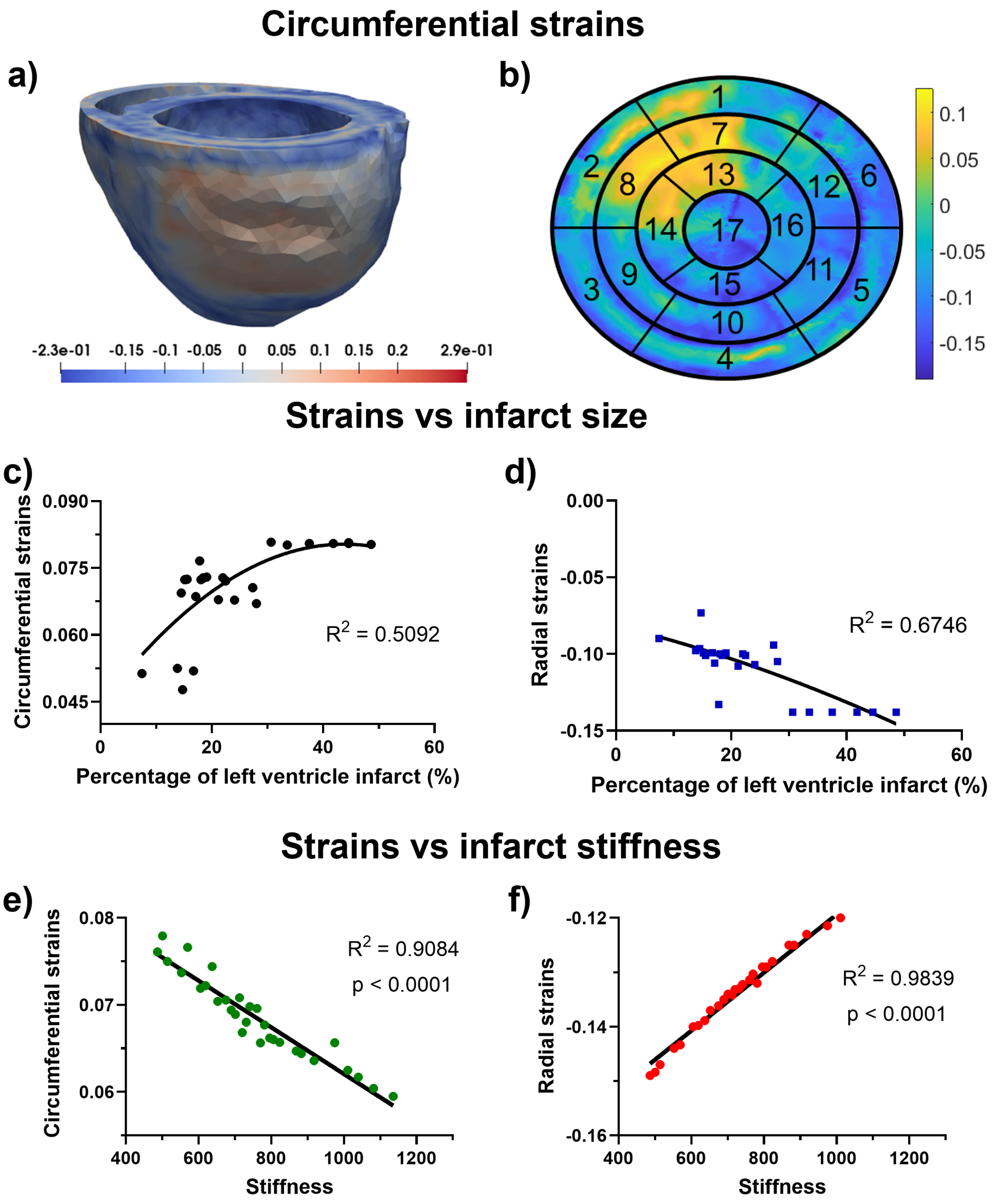


**Supplementary Fig. 1.** a) Representative in-silico circumferential strain distribution in a biventricular finite-element heart model, b) the corresponding circumferential strains of left ventricle (LV) using the American Heart Association segmentation, c) average circumferential strain of infarcted region in the LV versus the percentage of the infarcted region of the LV, d) average radial strain of infarcted region in the LV versus the percentage of the infarcted region of the LV, e) average circumferential strain of infarcted region in the LV versus stiffness of the LV, f) average radial strain of infarcted region in the LV versus stiffness of the LV.
